# Deciphering the molecular pathways of apoptosis using leaf extract of *Basella alba* against Ehrlichs Ascites Carcinoma (EAC) cell line in Swiss albino mice model

**DOI:** 10.1101/485078

**Authors:** Md. Shihabul Islam, Md. Sifat Rahi, Chowdhury Arif Jahangir, Md. Mahmudul Hasan, Salek Ahmed Sajib, Ariful Haque, Syed Rashel Kabir, Kazi Md. Faisal Hoque, Mohammad Ali Moni, Md. Abu Reza

## Abstract

A series of programmed cell death that plays a vital role in the exclusion of abnormal cells without any ruin into the surrounding neighboring cells is called Apoptosis. Generally, it occurs in multi-cellular organisms through an orderly and autonomously processes that is controlled by proper function of various genes. In the current studies, cell apoptosis in EAC cells treated with different fractions (BLP-01, BLP-02 and BLP-03) of leaf extract of *B. alba* was detected by conducting several bio-assays such as cell growth inhibition, fluorescence and optical microscopy, DNA fragmentation and PCR amplification etc. The results of these experiments indicate that the plant extracts are able to inhibit cell growth significantly where morphological features of apoptosis were appeared under both fluorescence and optical microscope. The PCR amplification results showed that the leaf extracts of *B. alba* were able to cause EAC cell apoptosis in both the extrinsic and intrinsic pathway. An excellent figure of fragmented DNA was found in DNA fragmentation assay when the gel was observed under UV light which confirms the cell apoptosis. The current findings suggest that the samples of this experiment occupy fascinating competence to conduct cell apoptosis and become an ideal resource for cancer research as well as drugs development for cancer treatment.

## 1. Introduction

Apoptosis is a genetic based cellular process occurring in pathological and physiological conditions. In 1972, the term apoptosis was first used by Kerr, Wyllie, and Currie to describe the several modes of cell death [1, 2]. Apoptosis is responsible for the removal of redundant and undesirable cells to regulate the sound health. Generally, it takes place in the time of improvement and aging to keep up cell populations in tissue through the homeostatic process [3, 4]. But it is proved that defective apoptotic pathway plays a central role to develop a number of neural and immune related disorders and diseases in human especially different types of cancer [5].

The different morphological alterations of cells that happen at the period of apoptosis have been detected by using either a light microscope or electron microscope. Through the function of light microscope, cell contractions are easily detected in the early stage of apoptosis and these are considered as the most important features of apoptosis [5, 6]. However, several physical changes of cells including membrane blebbing, apoptotic bodies, loss of membrane integrity, cytoplasmic modifications and so on are seen in late apoptotic phase [7].

Usually, a number of different biochemical alternations are detected in apoptotic cells namely activation of Caspases, cleavage of DNA, recognition of phagocytes, breakage of proteins, etc. [8]. The Caspases are the most important protein that generally exists in maximum cells in its pro-enzymatic form. The active Caspases accelerate the programmed cell death by initiating a protease cascade which expands the apoptosis signaling [9]. Similarly, they promote the protease and DNAse proteins activation which have a leading contribution to split the nuclear DNA, several cellular proteins, nuclear scaffolds, cytoskeletons and so on [10]. Researches indicated that, the recognition of phagocytes is considered as a crucial biochemical properties of apoptotic cells [11].

Mainly, there are two apoptotic pathways namely intrinsic pathway and extrinsic pathway [12, 13]. The intrinsic pathway is also known as non-receptor based pathway as well as mitochondrial apoptotic pathway which is initiated by intra-cellular stimuli such as radiations, toxic substances, hyperthermia, viral infections, hypoxia, free radicals etc. [5]. These stimuli activate some kinds of protein like cytochrome C that can promote to form an apoptosome by binding with Caspase-9. In this way, the action of this apoptosome inhibits the IAP (Inhibitors of Apoptotic Proteins) which accelerate the cell death as well as apoptosis [14, 15, 16]. Moreover, endonucleases G have vital contribution to cause DNA fragmentation and chromatin condensation [17]. On the other hand, the extrinsic pathway is also called as receptor mediated pathway in which several trans-membrane receptors are involved to start the apoptotic process. Some common receptors for the extrinsic pathway are Fas, TNF (tumor necrosis factor), Apo2L, Apo3L and etc. [18]. These receptors do not work in the absence of their analogous ligands. The interaction between receptors and ligands take place due to the presence of respective adaptor named death domains (TRADD, FADD). The receptor-adaptor- ligand complex activate the Caspase-8 that eventually agile the Caspase-3.

The active Caspase-3 sends death signal to intra-cellular signaling pathway from cell surface and initiates cell apoptosis [19, 20].

Apoptosis is very essential in different purposes as like as (i) to protect the body tissue from pathogen-invaded cells, (ii) to discard inflammatory cells, (iii) to knock out self invasive immune cells, (iv) to renovate the adult cells and so on [21, 22]. But defective apoptotic process can be initiated various human diseases like cancer. Cancer is a group of diseases involving in uncontrolled cell growth. The suppression of apoptotic process has a potential role in the cancer as well as tumor development [23, 24]. Though cancer is one of the most destructing abnormalities of human body in recent time but no effective treatment system is developed until today [25]. Hence, the scientists are trying to find out an efficient treatment system of cancer. An artificial induced apoptotic process may be beneficial for cancer treatment and in this case the natural resources are more suitable than synthetic one. Because it is proved that the synthetic agents can kill cancer cells alongside with normal cells [26]. Plants are the most important natural sources that contain several types of bio-active components which demonstrate abundant biological features with large applications in drinks, foods, perfumes, sanitary, cosmetics and medicine industries. Besides these, they have an important role in prevention of numerous diseases and disorders in human [27, 28]. For these reasons, the current investigation was set up to study the apoptotic activity of a plant source named *Basella alba* at molecular level on Swiss albino mice model bearing EAC cell line.

## 2. Materials and Methods

### 2.1 Collection and Preparation of Plant Materials

The fresh and green leaf of *B. alba* were gathered from the botanical garden (latitude: 24.374609 and longitude: 88.635472) of University of Rajshahi which is belonged to the Department of Botany and washed with distilled water properly. The collected plant is identified by Dr. A. H. M Mahbubur Rahman, Professor of Taxonomy, Department of Botany, University of Rajshahi, Rajshahi-6205, Bangladesh. The washed fresh leaves were then dried and made fine powder which was dissolved in absolute methanol eventually. Finally, the methanolic solution of leaves was filtered and lyophilized for further used.

### 2.2 Dose Preparation

The methanolic extract was separated through gel filtration chromatography by using LH-20 matrix. The gel filtration chromatography is also known as molecular sieve chromatography and separate the molecule according to their molecular weight and size [29, 30]. At first, the lyophilized samples were dissolved in methanol and filtered by gel filtration column into separate tubes to take optical density (OD) at 280 nm using spectrophotometer (GENESYS 10S UV-Vis). Based on their OD, the separated solutions were divided into three different pools like *B. alba* Leaf Pool-1= BLP-01, *B. alba* Leaf Pool-2= BLP-02, and *B. alba* Leaf Pool-3= BLP-03. The collected solutions of different pools were then kept into freeze dryer to dry for further use. Finally, the dried extracts were dissolved in 2% DMSO (Dimethyl Sulphoxide) at 1mg/ml concentration to prepare a stock solution. Each mouse of different group was treated with BLP-01, BLP-02 and BLP-03 as 12.0, 40.0 and 20.0 mg/Kg/day concentration respectively.

### 2.3 Experimental Animal and Cell line

The Swiss albino mice weighted approximately 25± 2.0 g were taken as an experimental animal for this study. These animals were collected from the animal house of Molecular Biology and Protein Science Laboratory, Department of Genetic Engineering and Biotechnology, University of Rajshahi, Bangladesh. The mice were nurtured in plastic cages with the proper environment (temperature 25±2.0^0^C and 12±1 hour dark/light cycle) and fed with rat pellet having standard nutrients. Thirty (30) mice were taken for this investigation which divided into 5 groups (each group consists of 6 animal) and marked as Group-01(for control), Group-02 (for BLP-01 treated), Group-03 (for BLP-02 treated), Group-04 (for BLP-03 treated), Group-05 (for standard drug Bleomycin treated, used only for growth inhibition assay). In addition, the experimental cell line, Ehrlich ascites carcinoma (EAC) cells were provided by Protein and Enzyme Laboratory, Department of Biochemistry and Molecular Biology, University of Rajshahi, Bangladesh. The cells were maintained *in vivo* condition into the intra-peritoneal cavity of Swiss albino mice.

### 2.4 Chemicals and Reagents

Chemicals used in the current project was Saline (1% NaCl), Sodium Citrate (Anti-coagulant), Methanol (Merk, Germany), Ethanol (Merk, Germany), Isopropanol, Phosphate buffer saline (PBS), 4,6-diamidino-2-phenylindole (DAPI), TIANamp Genomic DNA kit (Tiangen, Beijing, China), RNAsimple Total RNA kit (Tiangen, Beijing, China), Agarose (Fluka, Sweden), Ethidium bromide, TBE buffer, Trypan blue (Thermo Scientific, USA), Bromophenol blue, LH-20 matrix (GE Healthcare, USA), MTT 3-(4,5-dimethylthiazol- 2-yl)-2,5-diphenyltetrazolium bromide solution. All the chemicals and reagents were of laboratory and analytical grade.

### 2.5 MTT colorimetric bio-assay

The MTT assay was used to identify biological efficiency as well as the cytotoxic activity of the leaf extract. Initially, the EAC cells were isolated from mice peritoneal cavity and washed with PBS (Phosphate Buffer Saline) properly. Then the cells were taken into artificial cell culture media and 200 *µ*l of this mixer was transferred into 96 well cell culture plate (Nalgene, USA). Next, 100 *µ*l experimental extracts (3 different concentrations) were added into the wells for serial dilution and kept the plate at 37^0^C for 24 hours into the CO_2_ incubator. Subsequently, the aliquot was removed from each well and 20 *µ*l of MTT dissolved in 180 *µ*l of PBS solution were added into each well and the plate was kept for overnight incubation. After that, the aliquot was again discarded and added 200 *µ*l of acidic isopropanol solution and incubated for 1 hour. Finally, the absorbencies were taken at 570 nm to make graphs and calculated the LC_50_ value from the regression line.

### 2.6 Cellular study of Apoptosis

This study was conducted through cell growth inhibition bioassay in both *in vivo* and *in vitro* condition. Cell growth inhibition assay is one of the most effective tools to determine the anti-proliferative activity of biological compounds as well as plant materials on established cancer cell line or primary tumor cells. It is widely known that the natural compounds which are able to prohibit cell growth that can initiate apoptosis of cancer cells. These plant resources show the chemo-preventive activity through apoptosis process and reduce the number of cancer cells when cells are treated by them [31]. So, it is said that the reduction of cancer cells takes place through the apoptosis process due to treat with natural anticancer agents.

#### 2.6.1. In vivo cell growth inhibition

To evaluate the cancer cells growth inhibition for *in vivo* condition, a common and simplest method was used which was described by Sur et al. 1994 [32]. For therapeutic evaluation, each mouse of all groups was inoculated as 1.76 10^6^ EAC cells per ml concentration using narrow syringe. After 24 hours of inoculation, treatments were started with BLP-1= 300 *µ*g/ml, BLP- 2= 1000 *µ*g/ml and BLP-3 = 500 *µ*g/ml for each mouse and continued for six days. On 7^*th*^, days, the EAC cells were collected from the intra-peritoneal cavity of mice and diluted with normal saline water (1% NaCl) to prepare different slides with trypan blue stain for counting viable cells. The number of viable cells was counted by hemocytometer using equation-01 and the frequency of cell growth inhibition was calculated by using equation-02.

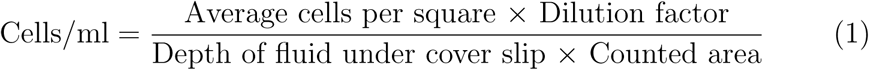

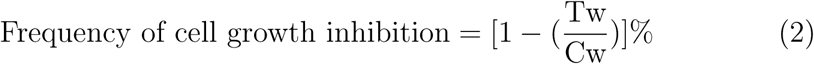

(Where Tw =Mean number of EAC cells for treated group and Cw = Mean number of EAC cells for control group mice).

#### 2.6.2. In vitro cell growth inhibition

To determine the EAC cell growth inhibition for *in vitro* condition, the experimental cells were cultured in artificial cell culture medium. For this study, first collected EAC cells were washed with PBS and diluted with 1% saline water for further uses. The cell culture medium was then prepared with proper nutrient and 10 ml of this culture medium was taken in several culture vessels where added 100 ml of EAC cells into each culture vessels eventually. After that, the cells were treated with experimental extracts at the similar doses of *in vivo* condition. Finally, the cultured cells were taken after 24 hours of treatment and prepared different slides for cell counting. The frequency of cell growth inhibition was estimated by equation-02.

### 2.7 Morphological examination of Apoptosis

A fluorescence microscope (Olympus iX71, Korea) was used to observe the morphological features of EAC cells for both control and treated cells. To perform this work, firstly, the cells were isolated and washed with PBS for 2-3 times after six days of treatment. Then, the cells were incubated with DAPI stain solution at 37^0^C for 20 min for staining. Subsequently, these cells were rewashed with PBS and prepare different slides. Finally, these slides were observed under light and fluorescence microscope and counted the apoptotic cells per slide.

### 2.8 Molecular Analysis of Apoptosis

When the morphological features are not good enough to prove cell apoptosis then DNA fragmentation assay and gene expression assay can be hugely used. Besides these, a number of techniques including TEM (Transmission electron microscopy), TUNEL etc. are applied which can give the confirm results for apoptosis [33, 34]. In the present study, the DNA fragmentation assay and gene expression assay (using different apoptosis related genes) were conducted to ensure cell apoptosis.

#### 2.8.1. DNA Fragmentation assay

To check the DNA breakage (an important feature of apoptosis) in this experiment, a reliable and easy technique named DNA fragmentation assay was used which first introduced by Andrew Wyllie AH, et al. in 1980 [35, 36]. In brief, firstly the total genomic DNA was isolated from both treated and control mice cells by using TIANamp Genomic DNA kit (Tiangen, Beijing, China) according to the manufacturers procedure. Subsequently, the isolated DNA samples were mixed with loading dye and run on a gel electrophoresis system for 50-60 minutes at 100 volts. Lastly, the gels were observed under UV light through a gel documentation system (Protein simple, Alpha imager mini, USA).

#### 2.8.2. Gene expression analysis

In the current study, the expression pattern of a number of genes which are related to cell apoptosis for example Fas, TNF-*α*, NF*κ*B, Bcl-2, Bcl-X, Bax, P-53, PARP-1, Cas-3, 8, 9, Cyto-C were detected through polymerase chain reaction (PCR) amplification technique. For this, initially the total RNA were extracted from both control and treated mice cells using RNAsample Total RNA kit (Tiangen, Beijing, China) by following company’s instructions and check the quality and quantity by nano drop (Thermo scientific) technique. After that, the isolated RNA was reverse transcribed to produce cDNA through RT-PCR method by following standard protocol. By using this cDNA the gene amplification was conducted with the help of 12 respective primers which sequence and annealing temperature (Tm) are shown in Table 1. Subsequently, the amplified PCR products were visualized under UV light using agarose gel electrophoresis system. 1kb plus DNA ladder (Tiangen, Beijing, China) was used as DNA marker. Finally, the expression levels of amplified genes were determined through Gel Analyzer-2010 software.

**Table 1.**
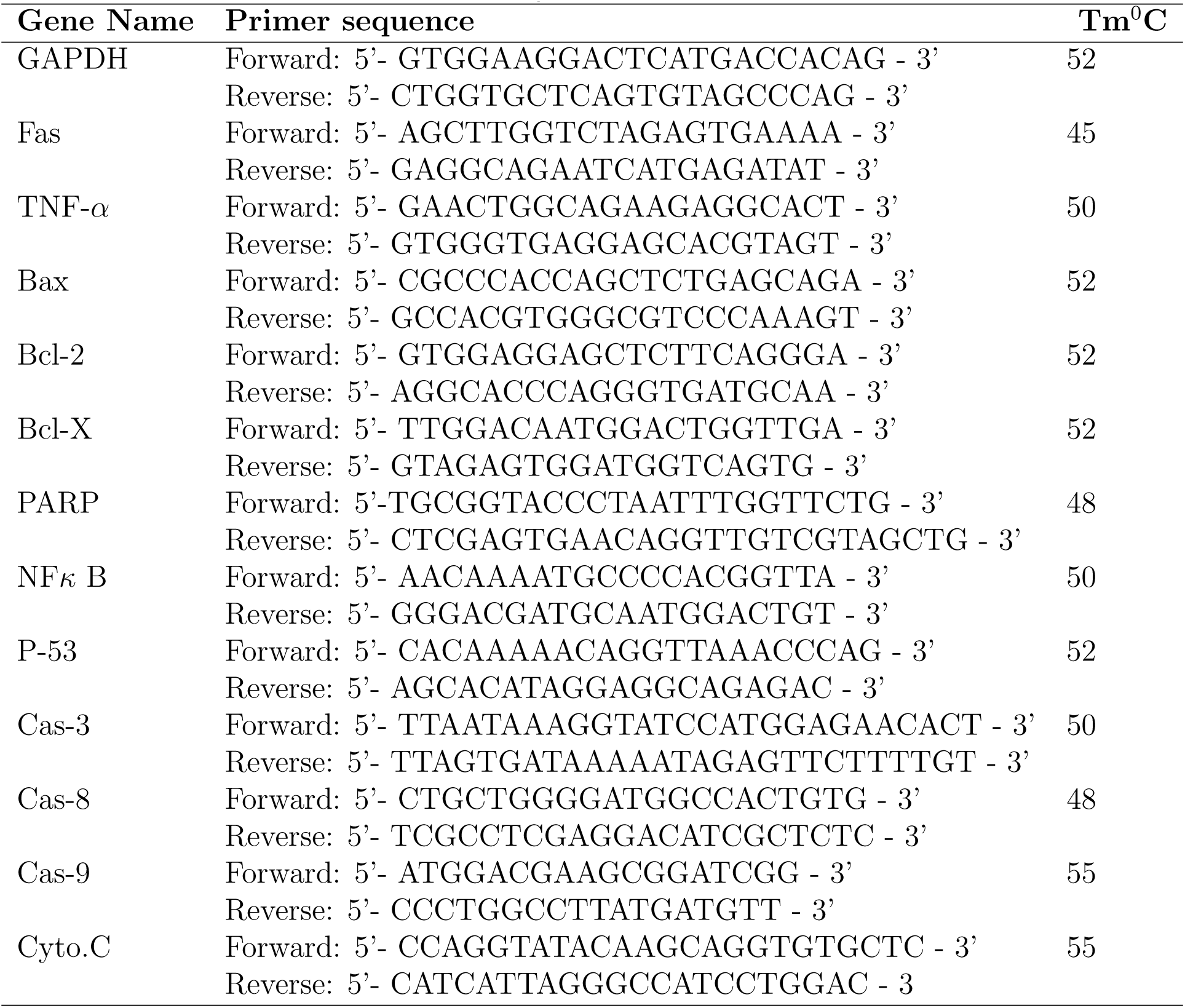
The sequence of primers used for PCR amplification with their respective annealing temperature

### 2.9 Statistical Test

Every statistical test was performed for thrice in time. All data are represented as mean±SD. The SPSS-16 software was used to test the significant of all the results through one-way ANOVA followed by Dunnett Post hoc test compare with control. Significance levels were set up at 5% (P*>*\*0.05), 1% (P*>*\*\*0.01) and 0.1% (P*>*\*\*\*0.001). For the statistical and graphical presentation of data, Microsoft Excel 2007 was used.

## 3. Results

### 3.1 MTT bio-assay

In MTT assay, BLP-01, BLP-02, and BLP-03 showed significant activity which is the indication that they are biologically potent. The 50% lethal concentration, (LC_50_) values of each test sample were calculated from the corresponding regression equation and the LC_50_ values of BLP-01, BLP-02 and BLP-03 were calculated as 308.66±5.28 *µ*g/ml, 1107.31±7.46 *µ*g/ml and 415.82±6.34 *µ*g/ml respectively, that is shown in Fig. 1(A). The BLP-01 showed the best bio-activity than BLP-02 and BLP-03 when compared with each other, exposed in Fig. 1(B). This result was very helpful to prepare the dose concentrations for treatment.

**Figure 1:**
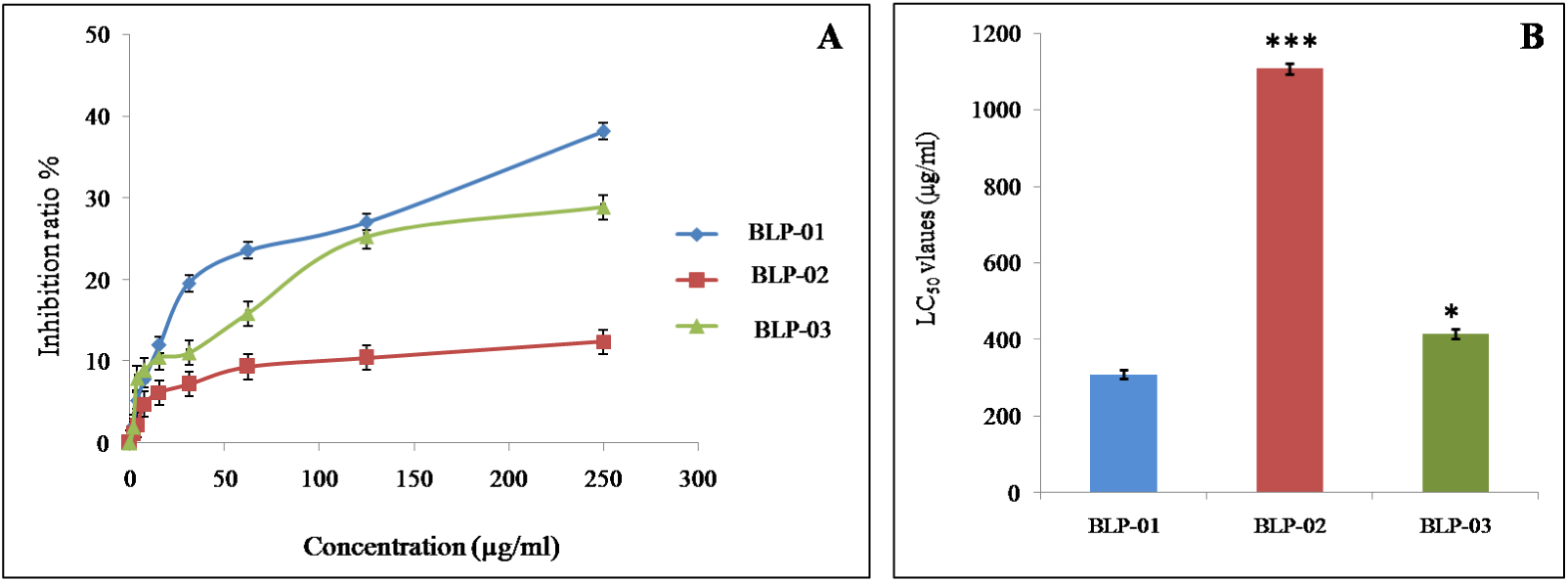
The graphical representation of MTT assay results of *B. alba* leaf extracts. (A) Inhibition frequency of BLP-01, BLP-02 and BLP-03 samples. (B) Comparison of LC50 values of three different extracts. Each value is presented as mean±SD (n=3) and significance was set at as 5% = P*<*0.05 (*), 1% = P*<*0.01 (**) and 0.1% = P*<*0.001 (***) when comparing with each another.

### 3.2 Cellular study of Apoptosis

The results of cell growth inhibition due to treatment with BLP-01, BLP- 02 and BLP-03 *in vivo* and *in vitro* condition were presented in Fig. 2(A) and Fig. 2(B) respectively. In case of *in vivo* situation, the proportions of cell growth inhibition of BLP-01, BLP-02 and BLP-03 treatment materials were measured as 71.22±2.54%, 51.39±3.61% and 45.53±2.91% respectively where inhibition percentage of Standard (Bleomycin) was 81.22±2.22%. On the other hand, the inhibition frequencies of BLP-01, BLP-02, and BLP-03 and standard *in vitro* condition were calculated as 46.85±1.62 %, 33.18±2.17%, 39.52±2.89% and 78.56±3.24% respectively. The hemocytometric counting of EAC cells demonstrated that the number of live cells declined due to treatment with leaf extracts of *B. alba* significantly when compared with control for both *in vivo* and *in vitro* condition. Based on the data that represented in Fig. 2(A) and Fig. 2(B), the BLP-01 has the highest capability to inhibit EAC cell growth.

**Figure 2:**
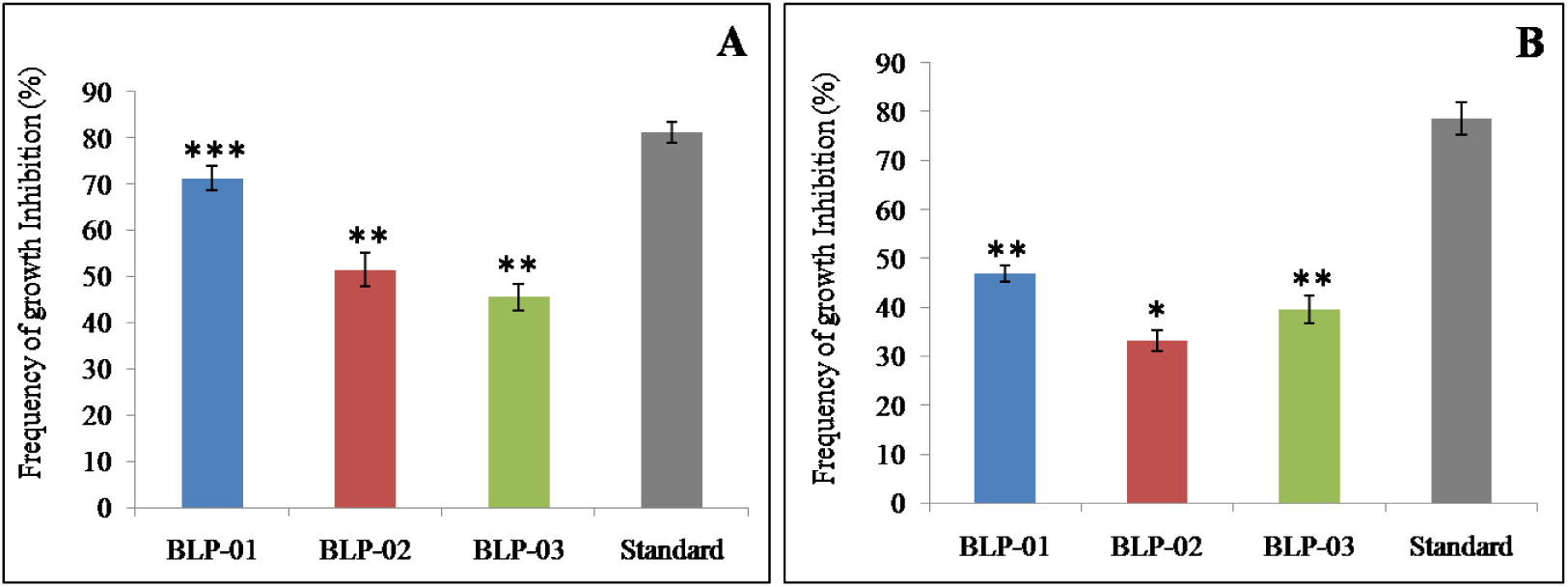
The cell growth inhibition of BLP-01, BLP-02 and BLP-03 extracts along with the standard (Bleomycin). (A) *In vivo* cell growth inhibition of experimental extracts (B) *In vitro* growth inhibition of EAC cell treated with leaf extracts of *B. alba*. A significant cell growth inhibition was observed in BLP-01, BLP-02 and BLP-03 treated mice when compared with the standard (Bleomycin) treated mice. All data are represented as mean±SD (where, n = 6). Significant tests are set up by comparing sample data with standard data and marked as 5% level = *P*<* 0.05, 1% level = **P*<*0.01 and 0.1% level=***P*<*0.001).

### 3.3 Morphological study of Apoptosis

The DAPI staining technique in this study reveals that, several nuclear alterations such as shrinked nucleus, condensed chromatin, nuclear breakage etc. were observed in all treated mice cells under fluorescence microscope where round and regular shaped nucleus were appeared in control mice that are shown in Fig. 3(A), 3(B), 3(C), and 3(D). Furthermore, in optical microscopic view, the cells were seemed to be regular shaped with normal cell membrane in control mice but the treated mice cells contained different structural changes including membrane blebbing, cell shrinkage, fragmented cell part, apoptotic bodies and so on, which are displayed in Fig. 3(E), 3(F), 3(G), and 3(H). All the abnormalities of EAC cells prove the performing of cell apoptosis treated by leaf extract of *B. alba*. The average number of apoptotic cells per slide for control, BLP-01, BLP-02 and BLP-03 treated mice were calculated as 4±1, 28±3, 16±2 and 10±2 respectively. The BLP-01 treated mice showed the significant number of apoptotic cells when compared with control which are expressed in Fig. 4.

**Figure 3:**
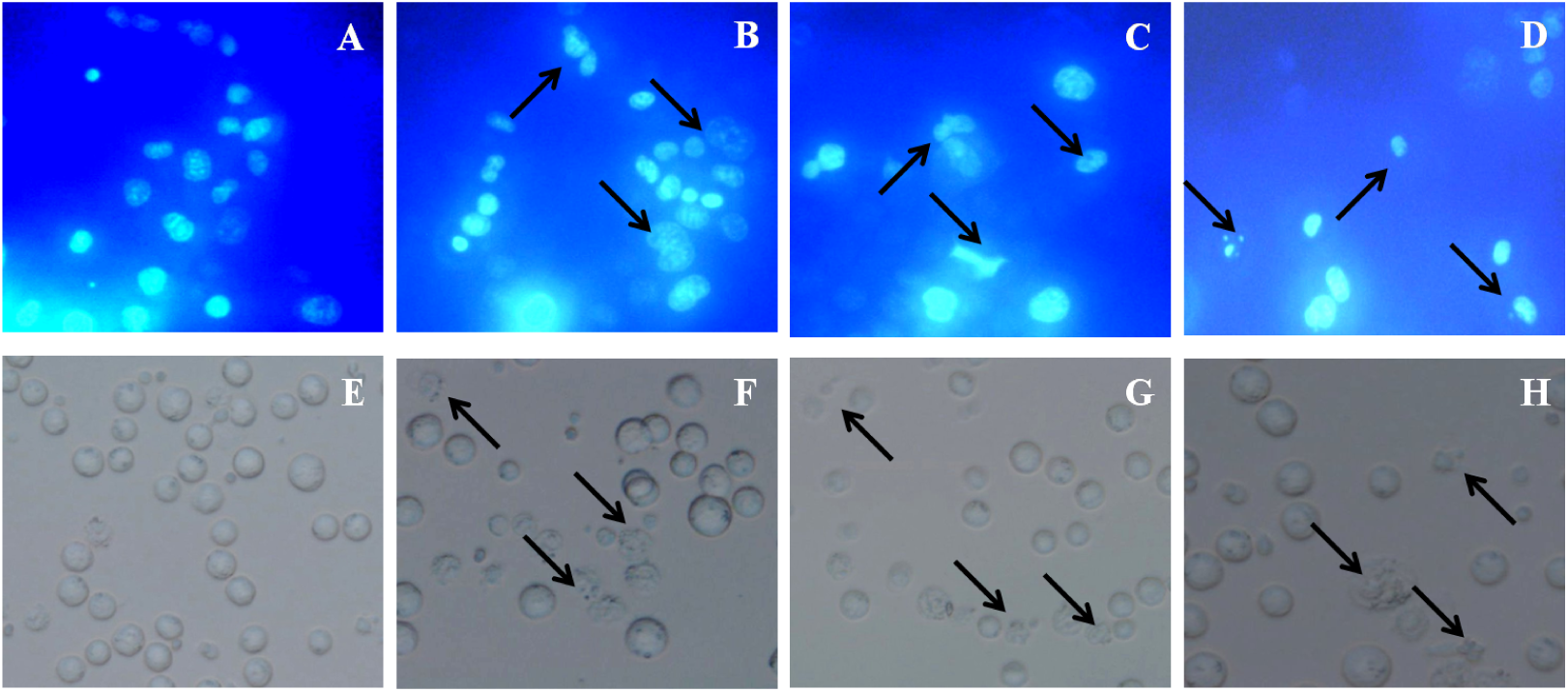
Determination of apoptotic cells by using DAPI staining technique for both control and treated mice after six days of treatment. The Fig. 3 (A, B, C, D) express as fluorescence microscopic observation and the Fig. 3 (E, F, G, H) represent as optical microscopic view of EAC cells for control, BLP-01, BLP-02 and BLP-03 treated mice respectively. Significant apoptotic features were seen in all treated EAC cells which are indicated by black arrows.

**Figure 4:**
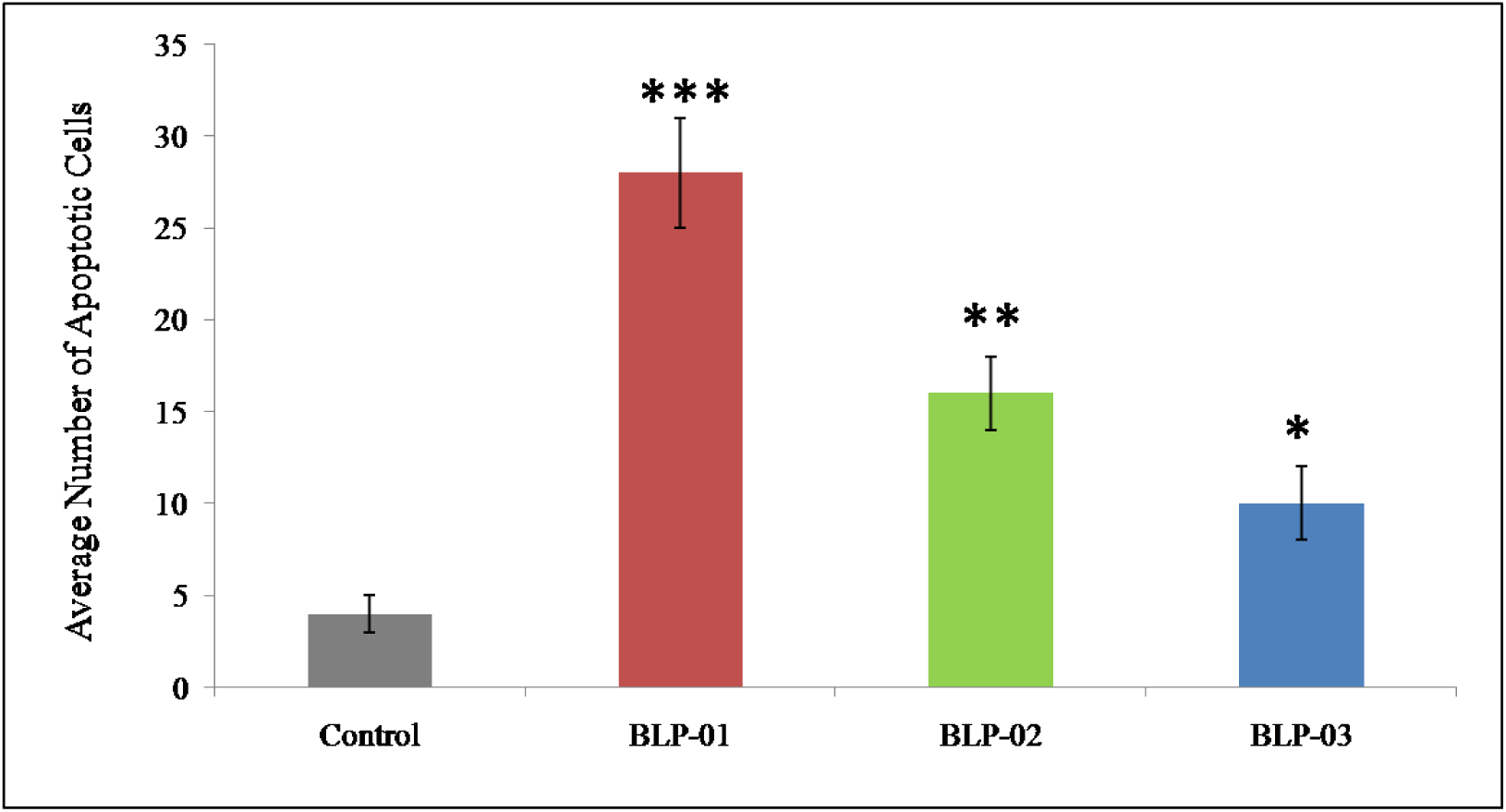
Average number of apoptotic cells per slide counted in both control and treated mice. Significant number of apoptotic cells was found in treated group mice in comparison with control. The result is presented as mean±SD (n=3). Significance was set at (*P*<*0.1, **P*<*0.01, and ***P*<*0.001) with respect to control.

### 3.4 Molecular Analysis of Apoptosis

For molecular analysis of apoptosis in EAC cells, two common techniques were used namely DNA fragmentation assay and gene amplification analysis. A high molecular weights band was appeared in control mice’s DNA lane on agarose gel that was deemed to be distinct and located at the top of the gel. But in case of treated mice’s DNA, fragmented or smeared DNA bands were seen at several locations on the gel that are presented in Fig. 5(A). This result confirms the apoptosis of EAC cells due to treat with investigated plant materials.

**Figure 5:**
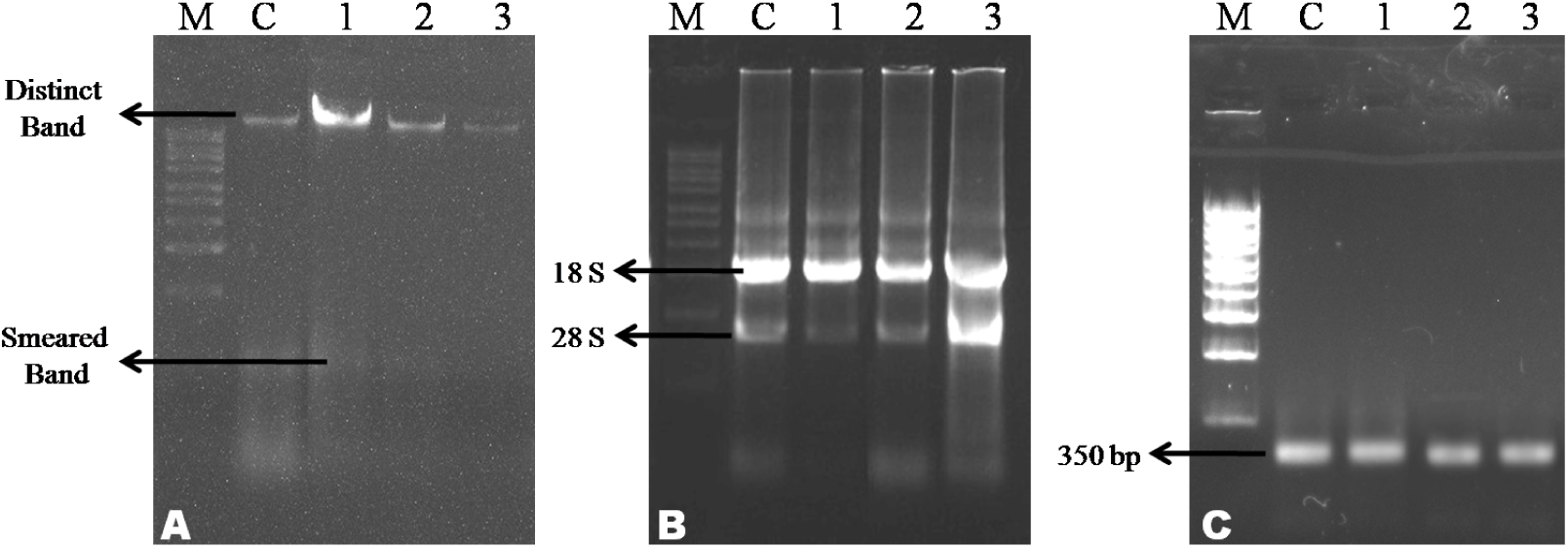
Molecular observation of apoptotic EAC cells for both control and treated mice. The lane M, C, 1, 2 and 3 are expressed as molecular marker, control, BLP-01, BLP-02 and BLP-03 respectively. (A) The banding pattern of DNA for treated mices DNA along with control (B) The agarose gel of RNA contained two different bands for both control and treated mice cells. (C) Gene expression assay by using GAPDH primer for treated and control mice.

For PCR amplification, the total RNA was isolated from both treated and control mice group and the Fig. 5(B) evident that the RNA was completely perfect that showed 28S and 18S RNA bands at the same level on gel. The expression pattern of GAPDH gene displayed in Fig. 5(C) confirms the successful conversion of cDNA from mRNA. By using this cDNA, PCR amplification was performed with 12 different primers associate with cell apoptosis. The banding pattern and expression level of these genes are shown in Fig. 6 and Fig. 7 respectively. The results of this molecular analysis indicated that experimental extracts are able to cause cell apoptosis through both intrinsic and extrinsic pathways. Based on the result of gene expression analysis, a proposed model was established that presented in Fig. 8 where both intrinsic and extrinsic pathway are shown.

**Figure 6:**
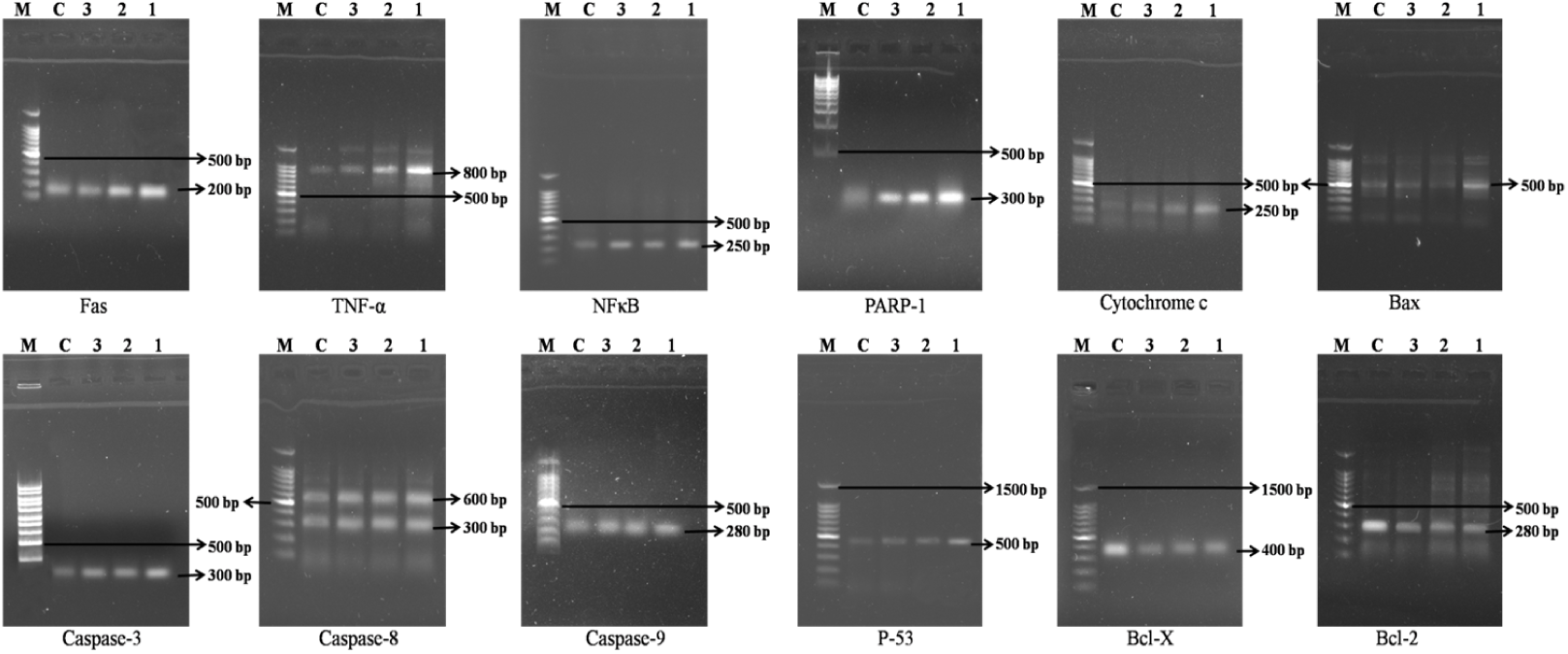
Agarose gel electrophoresis observation of 12 different genes related to cell apoptosis as well as cancer development for treated mice’s mRNA along with their respective controls. The lane M, C, 1, 2 and 3 express as molecular marker, control, BLP-01, BLP-02 and BLP-03 respectively.

**Figure 7:**
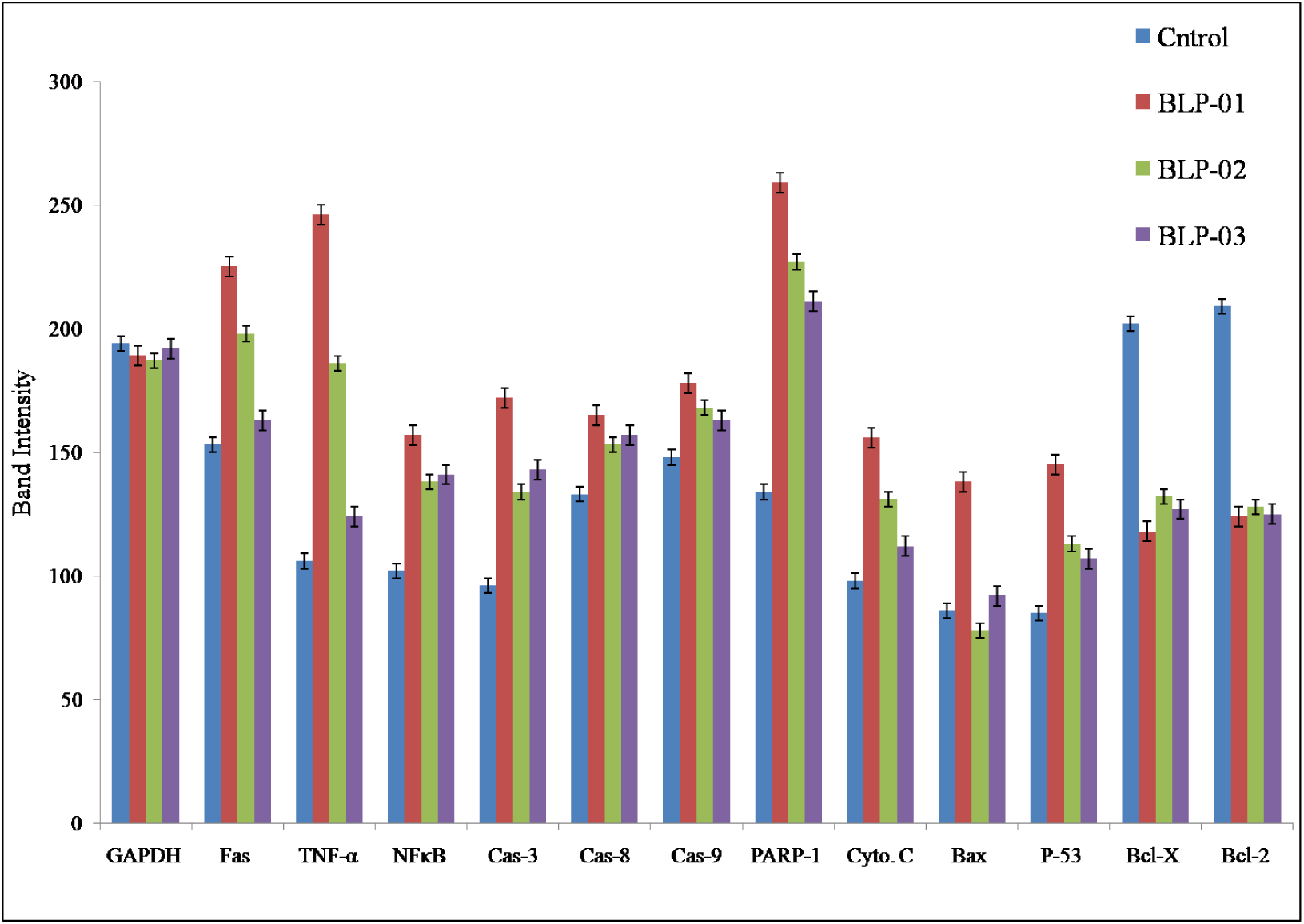
Relative observation of different gene expression levels for treated mice with their respective control based on their band intensity. Up regulated expression levels of Fas, TNF-*α*, NF*κ*B, Caspase-3, Caspase-8, Caspase-9, PARP-1, Cytochrome C, Bax and P-53 and down regulated of Bcl-X, Bcl-2 mRNA for treated mice groups than their respective controls confirm the EAC cell apoptosis at molecular level.

**Figure 8:**
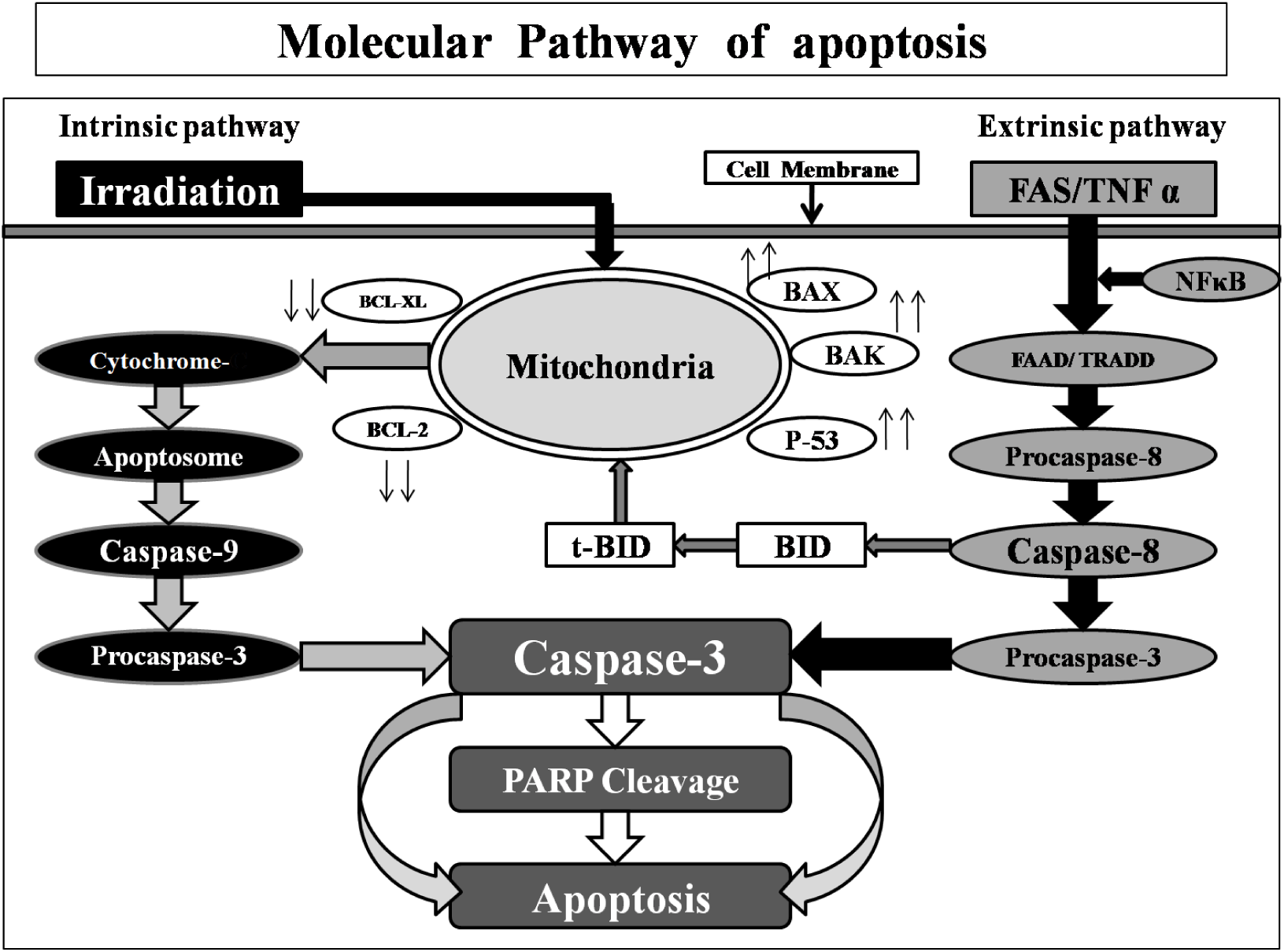
Proposed cell apoptosis model for both extrinsic and intrinsic pathway. This model was developed according to the results of gene expression analysis. The model showed, Cytochrome C, Caspase-9, Caspase-3, Bcl-X, Bcl-2, Bax, P-53 genes are involved in the intrinsic pathway and Fas, TNF-*α*, Caspase-8, Caspase-3, NF*κ*B, and PARP-1 genes are responsible for cell apoptosis through extrinsic pathway.

#### 3.4.1. PCR amplification of genes related to Intrinsic pathway of apoptosis

The intrinsic pathway of apoptosis is also known as mitochondria mediated pathway and a number of genes like Bcl-2, Bcl-X, Bcl-XL, P-53, Bax, Bak, Bid, Caspase-9, Caspase-3, cytochrome C etc. are responsible for this [37]. In the present study, up-regulation took place in P-53, Bax, Cas-9, Cas-3, cytochrome C genes and down regulation in Bcl-2, Bcl-X genes that are visualized in Fig. 6. The banding pattern and expression level of these discussed genes support the cell apoptosis through mitochondria mediated pathway.

#### 3.4.2. PCR amplification of genes associated to Extrinsic apoptosis pathway

The Fas, TNF-*α*, Caspase-8, NF*κ*B, Caspase-3, PARP-1 etc. genes are responsible for extrinsic pathway as well as receptor mediated pathway of apoptosis [38, 39]. The PCR amplification results in this experiment demonstrated the up-regulation of these genes in respect to their corresponding controls which are shown in Fig. 6. This result also proved the cell apoptosis which happened directly or by cleaving PARP-1 gene through extrinsic pathway.

### 3.5 Evaluation of gene expression level

The experimental gene expression levels were carried out based on their relative band intensity by using Gelanalyzer-2010 software. The results of the expression level for 12 different genes that were used in this study (mean while the GAPDH was used as standard) were presented in Fig. 7. According to these results, an equal expression level was seen in GAPDH gene for both control and treated mice where Bcl-2 and Bcl-X showed comparatively lower expression for treated mice than their respective control mice. But in the rest 10 genes, higher expression levels were found in treated mice than their corresponding controls.

## 4. Discussion

Apoptosis is a natural cell death process that is characterized by definite morphological features. It is established that the apoptotic process plays a central role in normal cell turns over, proper growth of immune system and chemical based cell death etc. But it’s deficiency is considered as a key factor of some human abnormalities including several types of cancer [3, 5]. It is also seemed that artificially induced apoptotic process by using natural phytochemicals reduces the growth of cancer cell. So that, the scientists and pharmaceutical companies become agree to find out natural components with different bio-activity for combating cancer development through inducing cell apoptosis [2, 40].

*B. alba* is a common vegetable plant that contains different bio-active compounds. In the current study, the bio-activity of *B. alba* leaf extracts was tested by conducting MTT assay which detects the potentiality of plants or plant parts initially [41]. In this experiment the notable LC_50_ values (Fig. 1) of plant materials help to prepare the doses for treatment. The plant derived bio-active substances can cause the induction of programmed cell death as well as cell growth inhibition.

Cell growth inhibition assay is one of the most popular bio-assays that is used worldwide to evaluate the frequency of viable cells in a collected sample using hemocytometer through trypan blue exclusion technique [42]. The leaf extracts of *B. alba* had an ability to inhibit the EAC cell growth due to the presence of several bio-active components. The experimental leaf extracts prohibited the cancer cell proliferation significantly in both *in vivo* and *in vitro* condition which is shown in Fig. 2(A) and 2(B) respectively. The BLP-01 showed (*in vivo* condition) the highest (71.22±2.54%) inhibitory activity than BLP-02 and BLP-03 where the standard Bleomycin (anticancer drug) showed 81.22±2.22% growth inhibition of EAC cells.

The present experiment reported that the BLP-01, BLP-02 and BLP-03 could initiate not only the cell growth inhibition but also altered the morphological changes of EAC cells which are the primary indication of cell apoptosis. The unhealthy and non-essential cells are removed from the body except any damages of adjacent cells through this process. The inhibition of the apoptotic process offers the normal and healthy cells to grow unusually that initiate the tumor as well as cancer development subsequently [43]. However, the induction of apoptosis by artificial medium have a vital role in the prohibition of several kinds of cancer cell [44]. The DAPI staining technique exhibited the deflect morphological properties of EAC cells including cell contraction, chromosome coagulation, nuclear alteration, aggregation of apoptotic bodies and so on due to treat with BLP-01, BLP-02 and BLP-03 where normal and round shaped cells were seen in control mice, presented in Fig. 3. Moreover, relatively less amount of apoptotic cells were counted in treated group mice than control that is shown in Fig. 4.

Only morphological view of cells is not enough to describe apoptosis confirmation. Besides this, molecular analysis is required to be ensured of cell apoptosis. DNA fragmentation bio-assay is one of the most popular and common ways to insure cell apoptosis at molecular level [35]. The results of current study reported that the experimental plant samples cleave the large sequence of DNA that generated smeared or fragmented bands on the agarose gel that is displayed in Fig. 5(A).

In the molecular point of view, the intrinsic and extrinsic pathways are two established pathway of cell apoptosis. The intrinsic pathway that is also called non-receptor related way which initiates by interactions among different types of proteins controlled by anti-apoptotic and pro-apoptotic genes. The abnormal functions of these genes promote to create mitochondrial pores which enhance the cytochrome C protein to form apoptosome by binding with Apaf-1. Later, the apoptosome activates the Caspase-9 that help to take place the cell apoptosis by activating the Caspase-3 subsequently [45, 46]

In contrast, the extrinsic or receptor mediated pathway initiates by the interaction between death receptors (FasR, TNF-*α* R) and their corresponding ligands (Fas, TNF- *α*). These clusters bind with their respective death domain (FADD or TRADD) to promote the Caspase-8 activation. The active Caspase-8 activates the Caspase-3 which causes apoptosis either directly or cleaving PARP-1 gene through DNA degradation. In the present investigation, the down-regulation of Bcl-2 and Bcl-X genes was found where other rest 10 used genes showed up-regulation in treated mice’s mRNA in respect to their corresponding controls (presented in Fig. 6 and Fig. 7). This result is a strong evident for apoptosis of EAC cells through both intrinsic and extrinsic pathway. The model exposed in Fig. 8 illustrates that our experimental extracts inhibit the EAC cells proliferation as well as initiate cell apoptosis through not only intrinsic but also extrinsic pathway.

## 5. Conclusion

Cancer is one of the most life threatening diseases in the world existence with limited treatment system and it is crying need to look for an efficient way of getting rid from this dangerous disease using natural resources. The plant *B. alba* is a common natural source with different bio-activity. The present study demonstrated that the methanol leaf extracts of *B. alba* are able to reduce the EAC cell’s growth and change the cell’s shape as well. They rooted out the cancer cells by DNA degradation and apoptotic process. The molecular study ensured that the experimental plant materials have the capability to apoptosis of EAC cell through not only receptor mediated extrinsic pathway but also mitochondrial based intrinsic pathway. However, more investigations are required to detect the noble compounds that are really responsible for cancer cell death by apoptotic process.

## Ethical Clearance

This experimental work was authenticated by the authority of Institute of Biological Sciences, University of Rajshahi-6205, Bangladesh. The Institutional Animal, Medical Ethics, Bio-safety, and Bio-security Committee (IAMEBBC) accepted this work for Experimentations on Animal, Human, Microbes and Living Natural Sources. The memo no. of this experiment: 31/320- IAMEBBC/IBSc.

## Acknowledgment

This research work was funded by the Molecular Biology and Protein Science Laboratory, Department of Genetic Engineering and Biotechnology, University of Rajshahi- 6205, Bangladesh.

